# Exosome labeling by lipophilic dye PKH26 results in significant increase in vesicle size

**DOI:** 10.1101/532028

**Authors:** Mehdi Dehghani, Shannon M. Gulvin, Jonathan Flax, Thomas R. Gaborski

**Affiliations:** Department of Microsystems Engineering, Rochester Institute of Technology, Rochester, NY, United States; Department of Biomedical Engineering, Rochester Institute of Technology, Rochester, NY, United States; Department of Urology, University of Rochester Medical Center, Rochester, NY, United States; Department of Biomedical Engineering, University of Rochester, Rochester, NY, United States

**Keywords:** EVs, PKH, CFSE, Size, Nanoparticle Tracking Analysis, Cellular Uptake, Biodistribution, Fluorescent Labelling, Lipophilic dye

## Abstract

Extracellular vesicles (EVs) are membrane vesicles secreted by cells and distributed widely in all biofluids. EVs can modulate the biological activities of cells in a paracrine or endocrine manner, in part by transferring their content, such as miRNA, following uptake in recipient cells. Fluorescent labelling of EVs is a commonly used technique for understanding their cellular targeting and biodistribution. Lipophilic fluorescent dyes such as those in the PKH family have been widely used for EV labelling. One concern with the use of lipophilic dyes is an increase in the EV size. This size shift alone may undermine the validity of EVs tracing studies as small changes in the size of inorganic nanoparticles are known to affect their cellular uptake and biodistribution. Here, the possibility of minimizing the size shift of PKH labelled EVs was systematically studied by changing the labelling condition. Unfortunately, the size shift towards larger particles was observed in all the PKH labelling conditions, including those where the labelled EVs were below the fluorescent detection limit. As opposed to lipophilic dyes, no significant shift in the size of labelled EVs was detected with protein binding dyes. Since the size shifts identified in all the PKH labelling conditions are likely to affect the cellular uptake and biodistribution, PKH may not be a reliable technique for EVs tracking.

## Introduction

EVs are small-membrane bound vesicles (30–150 nm in diameter) secreted by all cell types examined and can be found in almost all biofluids including blood, breast milk, urine, saliva and even in cell culture media [1,2]. EVs mediate cell-cell communication by exchanging proteins, DNA, RNA and lipids between donor and recipient cells and activating signaling pathways in target cells via receptor ligand interaction [3,4]. They have been shown to play a role in regulating both physiologic and pathologic processes including immune regulation and cancer [5]. EVs secreted by cells into the extracellular environment have been shown to be internalized through different routes and mechanisms including fusion with the plasma membrane and a range of endocytic pathways such as receptor-mediated endocytosis, phagocytosis, lipid raft-dependent endocytosis and micropinocytosis [6–9]. EVs target a wide range of recipient cells such as dendritic cell [10], macrophage [11], dermal fibroblast [12], endothelial and myocardial cells [13].

Studies that examine EV uptake into target cells and *in vivo* biodistribution have utilized a range of EV labelling and tracking approaches to follow EV fate [14]. The most common technique for studying EV biodistribution and target cell interaction involves labelling of EVs with fluorescent dyes. EVs membranes have been stained using the fluorescent lipid membrane (lipophilic) dyes like PKH26 [15–17], PKH67 [18], DiI [19], and DiD [20]. The PKH family have been the widely used dyes in the lipophilic dyes class as they have a highly fluorescent polar head group and long aliphatic hydrocarbon tails which readily intercalates into any lipid structure leading to a long-term dye retention and stable fluorescent [15,20].

Despite the widespread use of lipophilic dyes (PKH) for labelling of EVs [10,12,16,21], recent studies have highlighted generation of artifacts such as formation of numerous nanoparticles which consist exclusively of micelles/aggregates of PKH, without EV content [22–24]. It was further shown that in terms of size, surface area and fluorescent intensity, the PKH nanoparticles cannot be distinguished from PKH labelled EVs and were taken up by astrocytes. This capacity for cell uptake of PKH nanoparticles may lead to false positive signals in EV tracking studies [24]. However, cyanine-based membrane probes, called Membright have been recently developed which do not form nanoparticles in contrast to commonly used PKH family [25,26].

Up to now, far too little attention has been paid to generation of larger species after lipophilic labelling of EVs. A lipophilic dialkylcarbocyanine tracer (DiI) was used to enhance clustering and aggregation of EVs, confirmed by electron microscopy and flow cytometry [27]. Furthermore, a FM lipophilic styryl dye was found to result in larger EVs after labelling characterized by flow cytometry [28]. Additionally, using Nanoparticle Tracking Analysis (NTA), size distribution of EVs before and after labelling with PKH dye was studied. PKH labelling of EVs showed generation of larger particles which was not mentioned by the authors in the study [23].

This larger PKH-labelled EV species may alter uptake into target cells as previous studies have shown that cells preferentially uptake smaller EVs [29]. However, the contribution of size versus EVs constituents in this uptake phenomena is unclear. In order to examine the impact of particle size alone, the size-dependent uptake of nanoparticles composed of inorganic materials including those made of polystyrene [30] and silica [31] has been studied. These studies have shown that the interaction and uptake of nanoparticles with living cells are sensitive to the size of nanoparticles regardless of their composition. Lower cellular uptake of nanoparticles was consistently observed with increasing nanoparticle size, possibly due to the increased energy required to take up the larger particles [14,32–34]. Furthermore, size affects the biodistribution of nanoparticles *in vivo*, for instance, more rapid accumulation of larger nanoparticles was observed in liver and spleen [35]. Large particles also tend to exhibit shorter circulation half-life, which may be due to the activation of the complement system and quick removal of large nanoparticles from blood [36].

Since PKH family has been widely used for labelling EVs [10,12,16,21], formation of larger species after labelling EVs with PKH may trigger abnormalities in their tissue distribution and cellular uptake in both *in vivo* and *in vitro* studies. The objective of this study was to investigate the possibility of minimizing the formation of larger labelled EV species. In order to minimize the formation of larger labelled EVs, the PKH to EVs ratio was systematically changed and the size of particles was determined using Nanoparticle Tracking Analysis (NTA). In all labelling conditions including where the labelled EVs were below the fluorescent detection level, EVs size was increased.

## Materials and Methods

### Fluorescent staining

Lyophilized urinary CD63, CD9, CD81 positive EVs (HansaBioMed, Estonia) were diluted in ultra-pure water to a protein concentration of 0.1 μg/mL, following the manufacturer’s instructions. Prior to staining, 1 μM of PKH26 (Red Fluorescent Cell linker for General Cell Membrane, Sigma-Aldrich) stock was added to 300 μL of diluent C and incubated at 37° C for 15 minutes. Then, 1 μL of EVs stock was added to PKH26 in diluent C, resulting in a sample with 0.3 μg/mL of EVs and 4 μM of PKH26. Other concentrations of EVs and PKH were made accordingly. CFSE (CellTrace™ CFSE Cell Proliferation Kit, Thermo Scientific Fisher) stock was made following the manufacture instructions by adding 18 μL of Dimethyl Sulfoxide (DMSO, Sigma-Aldrich) to the CFSE resulting in 5 mM stock. In order to stain EVs with CFSE, 1 μL of CFSE stock was added to 300 μL of PBS prior to staining and then 1 μL of EVs from 0.1 μg/mL EVs stock was added and incubated for 2 hours at 37° C as it was previously described [23]. In order to remove unbound dyes, the samples were ultracentrifuged (Optima MAX-XP, Beckman Coulter) at 100000 x g for 60 minutes at 4 °C (TLA 110 rotor with cleaning factor of 81).

### Nanoparticle tracking analysis (NTA)

For each run, 300 μL of the prepared samples was injected into the sample chamber of a NS300 instrument (NanoSight, Aumesbery, UK) with a green laser 532 nm. Seven measurements of the same sample were performed for 30 seconds. For the “Blur”, “Minimum expected particle size”, and “Minimal track lengths” the auto adjustment settings provided by software developer were used. The camera level (9-12) and detection threshold (2-6) were adjusted manually for each experiment as recommended by the manufacturer. For data capturing and analysis, the NTA analytical software (NanoSight NTA 3.2) was used. Briefly, from the recorded video, the mean square displacement of each detected particle was determined. Then, using the Stokes-Einstein equation, the diffusion coefficient and sphere-equivalent hydrodynamic radius were determined by the software.

### Fluorescent imaging and analysis

Fluorescent images were taken using Keyence BZ-X700 microscope (Keyence Corp. of America, MA, USA) with the same exposure time for all samples. The line scan analysis was conducted using NIH ImageJ.

## Results and Discussion

Lipophilic dyes such as the PKH family have been widely used to label a range of cell types such as mesenchymal stem cells [17,37] and tumor cells [38] in proliferation and migration studies [21,37]. Since EVs have a lipid bilayer structure similar to that of the cell’s plasma membrane, PKH dye family have been adapted for EV labelling. As noted above, Dominkus et al. [24] have recently shown that PKH labelling of EVs generates larger species. In order to confirm these findings, EVs were labelled with a similar PKH to EVs molar ratio and the resulting particles’ size was assessed using NTA. NTA is a technique for measuring the size and concentration of nanoparticles in suspension in real time based on tracking the light scattered from suspended particles undergoing Brownian motion [39–41]. As was seen previously [23,24], nanoparticles were present in PKH only controls and critically, the generation of large species, greater than that found in either the EVs or PKH only controls, was observed following labelling of EVs with PKH (Figure 1A and 1B). Additionally, the interaction of 100 nm polystyrene (PS) nanoparticles with PKH was studied as a control experiment. In contrast to EVs, PKH should not interact with PS nanoparticles and as expected, the formation of larger particles was not observed when 100 nm PS nanoparticles were added to the PKH sample (Figure 1C).

**Fig. 1.**
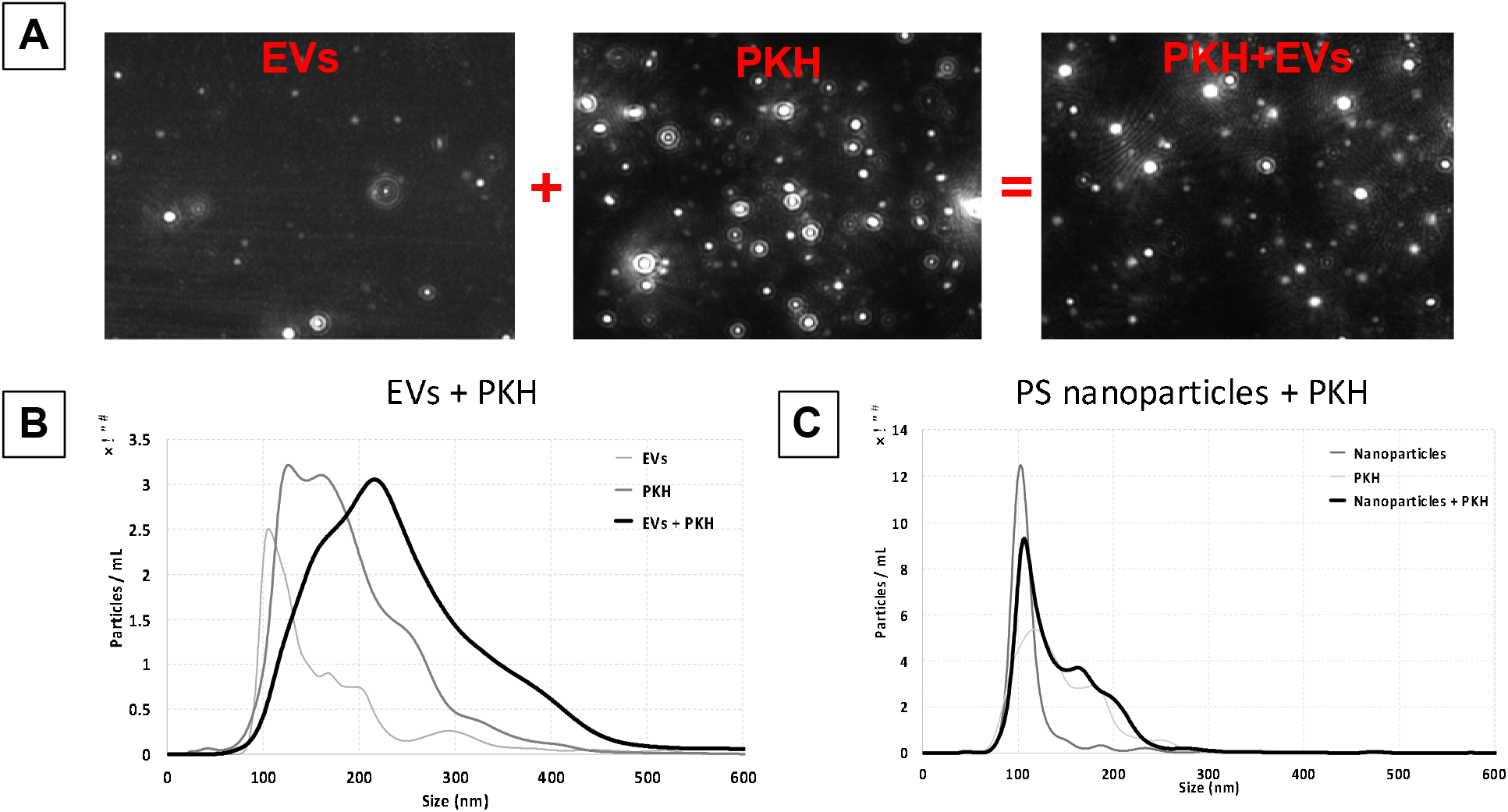
Size characterization of PKH EV labelling by Nanoparticles Tracking Analysis (NTA). A) NTA video frames of EV only (EVs) as control, PKH only (PKH) as control, and PKH-labelled EV (PKH+EVs). B) Size distribution of EV only control (EVs), PKH only control (PKH), and PKH-labelled EV (PKH+EVs) samples (n=7). C) Size distribution of PS nanoparticles only control, PKH only control, PS nanoparticles in PKH sample (n=7).

Possible mechanisms causing the observed size increase are the aggregation/fusion of nanoparticles with EVs or the intercalation of PKH molecules into EV membranes, all of which would result in the formation of larger species. This suggests the possibility of minimizing the size increase by reducing the PKH input level, while maintaining detectable fluorescent signal from PKH labelled EVs. To assess this possibility, the fluorescent detection range of PKH-labelled EVs samples was determined. PKH/EV ratios were adjusted by changing the concentration of PKH while holding the EVs concentration constant. The fluorescent level of these samples along with that of the PKH and EV only, and background only control groups was visualized using fluorescent microscopy and quantitated by sampling the cross-sectional fluorescent intensity of captured images (the line scan).

Initially, the same concentration of PKH and EVs was used as in figure 1. Weakly fluorescent features consistent with the presence of PKH nanoparticles in the PKH only control were observed by fluorescent imaging (Figure 2A). In comparison to the PKH only control, several brighter features were seen which are likely the larger particles formed after PKH EVs labelling (Figure 2B). The line scan of PKH-labelled EV samples showed higher fluorescent intensity compared to the background signal (Figure 2E). Line scan analysis of the PKH only control and PKH-labelled EV samples revealed a reduction in the baseline signal possibly as the result of floating PKH dyes interacting with EVs (Figure 2E). Furthermore, larger spikes in the PKH-labelled EV sample again suggesting the presence of larger labelled EVs (Figure 2E). In contrast, a 25 times reduction of PKH concentration led to a decrease in the fluorescent intensity to the same signal level as the background, as well as causing the loss of the fluorescently bright features observed at higher concentrations of PKH (Figure 2C-E).

**Fig. 2.**
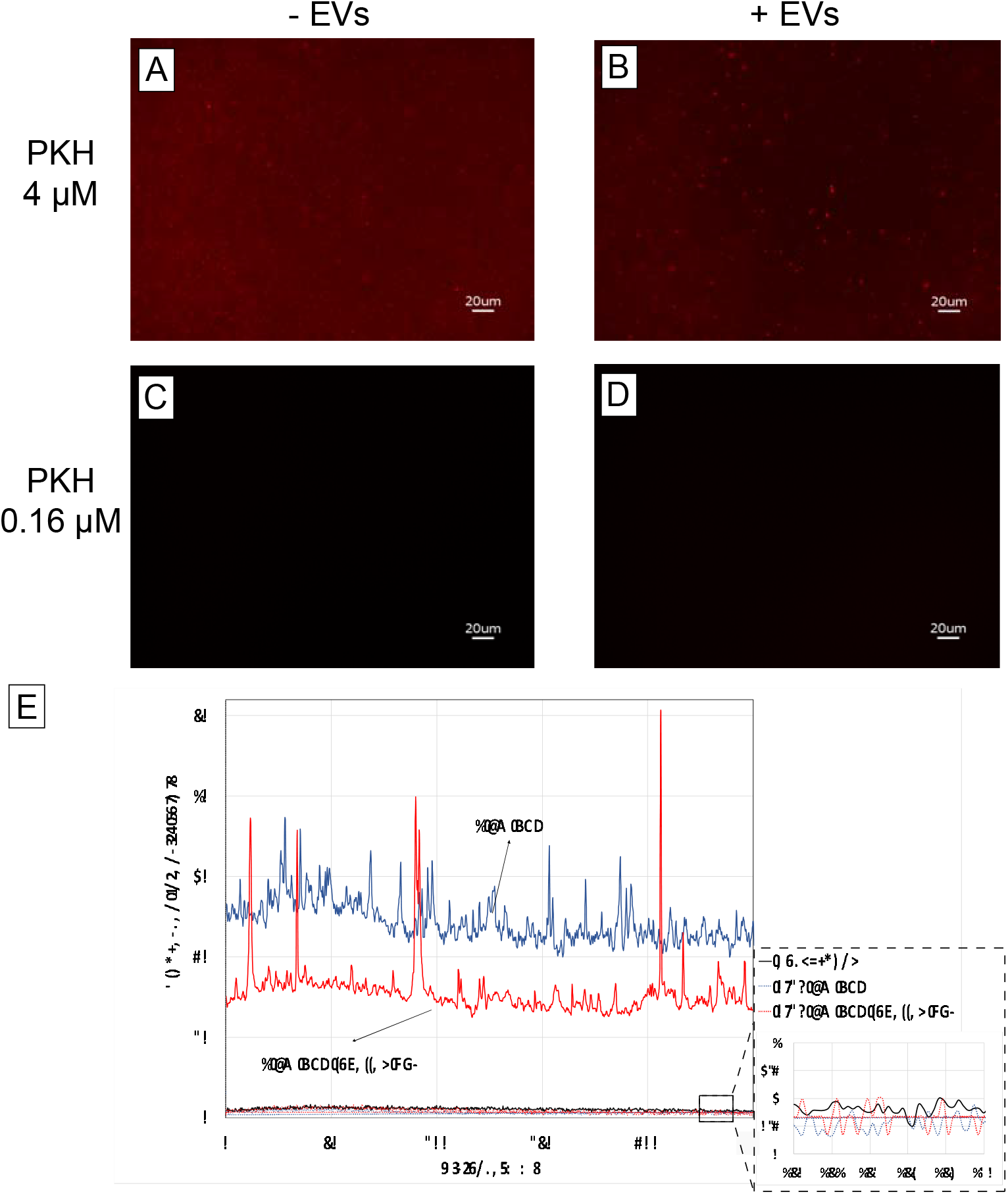
Determination of the fluorescent detection range. Fluorescent images of A) 4 μM PKH only control B) 4 μM PKH-labelled EVs C) 0.16 μM PKH only control D) 0.16 μM PKH-labelled EVs. E) Representative line plot analysis of fluorescent images (A-D). Inset is showing the line plot analysis of background, 0.16 μM PKH only control and 0.16 μM PKH-labelled EVs.

After determining the fluorescent detection range of PKH labelled EV samples, the effect of PKH concentration on the particles size distribution was explored. The concentration of PKH was systematically varied while holding EV concentration constant. Representative examples of particle size distribution measured by NTA for different concentration of PKH labelling of EVs can be seen in figure 3 (A – D). For all concentrations of PKH, NTA analysis of particle size distribution showed that EV labelling with PKH caused the formation of larger species relative to EVs and PKH alone and EV alone control groups (Figure 3D). Quantitative determination of NTA results was done by comparing the mode of the nanoparticle size (Figure 3E). Consistent with size distribution results, a shift in the mode towards larger particles was observed in all PKH-labelled EV samples (PKH+EVs) compared to the EV alone control. Furthermore, no size shift was observed when the suspension buffer (diluent C) was added to EVs in the absence of PKH confirming that PKH is the cause of the size shift.

**Fig. 3.**
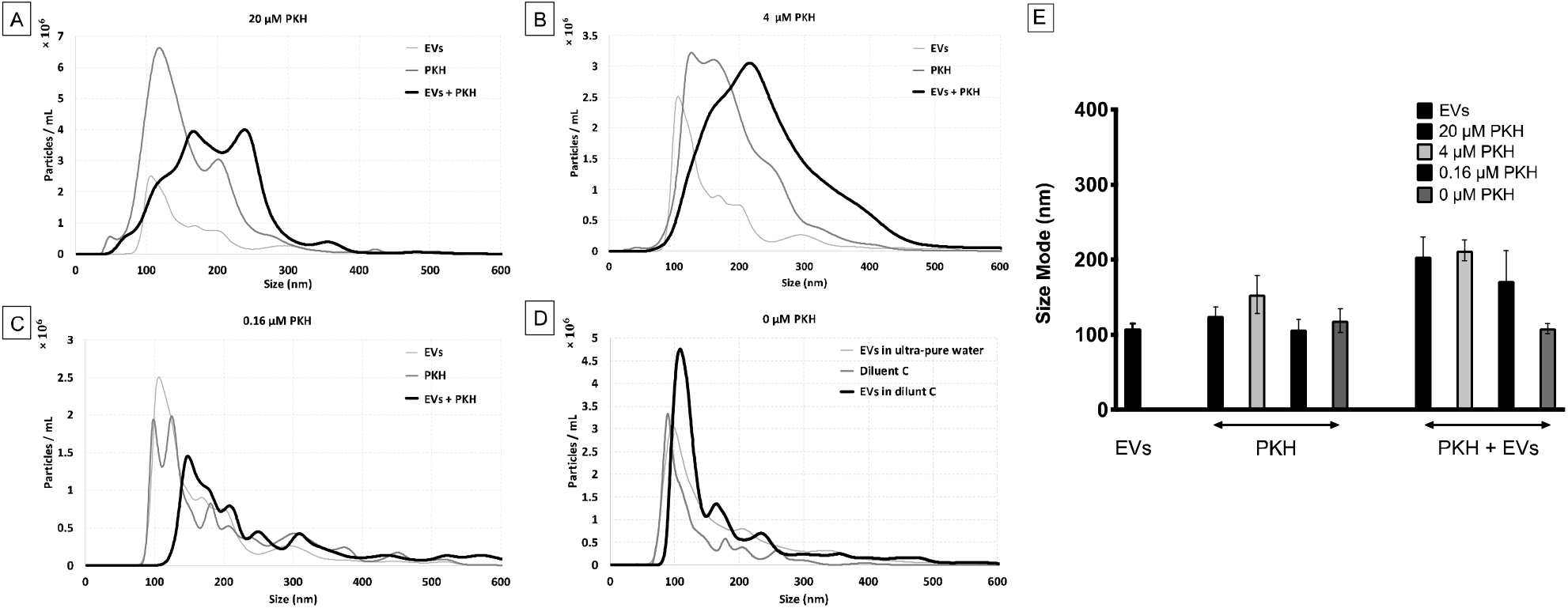
Effect of PKH concentration on the size distribution of particles in PKH-labelled EV samples by Nanoparticles Tracking Analysis (NTA). A-D) Size distribution of PKH-labelled EV samples with different PKH concentrations (20, 4, 0.16 and 0 μM respectively) with 0.3 μg/mL of EVs (n=7). E) Particles size mode for EV only control, PKH only controls, and PKH-labelled EV samples (Error bars represent the 95% confidence interval of the mean).

Taken together, labelling EVs with different concentrations of PKH, even for the PKH concentration below the level of fluorescent detection, showed the size distribution shift indicative of the generation of larger species. This finding suggests that minimizing the formation of larger species by reducing the concentration of PKH used for labelling EVs, while maintaining the fluorescent detectability may not be feasible.

Further confirmation of PKH induced larger species was done by varying the concentration of EVs while holding the PKH concentration constant. The PKH concentration used was the level shown to generate fluorescently detectable PKH-labelled EVs (Figure 2B). Representative examples of particles size distribution measured by NTA for different concentration of EVs labelled by PKH dyes can be seen in figure 4 (A – D). Additionally, quantitative determination of NTA results was conducted by comparing the mode of the nanoparticles size (Figure 4E). As expected, generation of larger species was observed by size distribution as well as the shift in the size mode regardless of the EVs concentration.

**Fig. 4.**
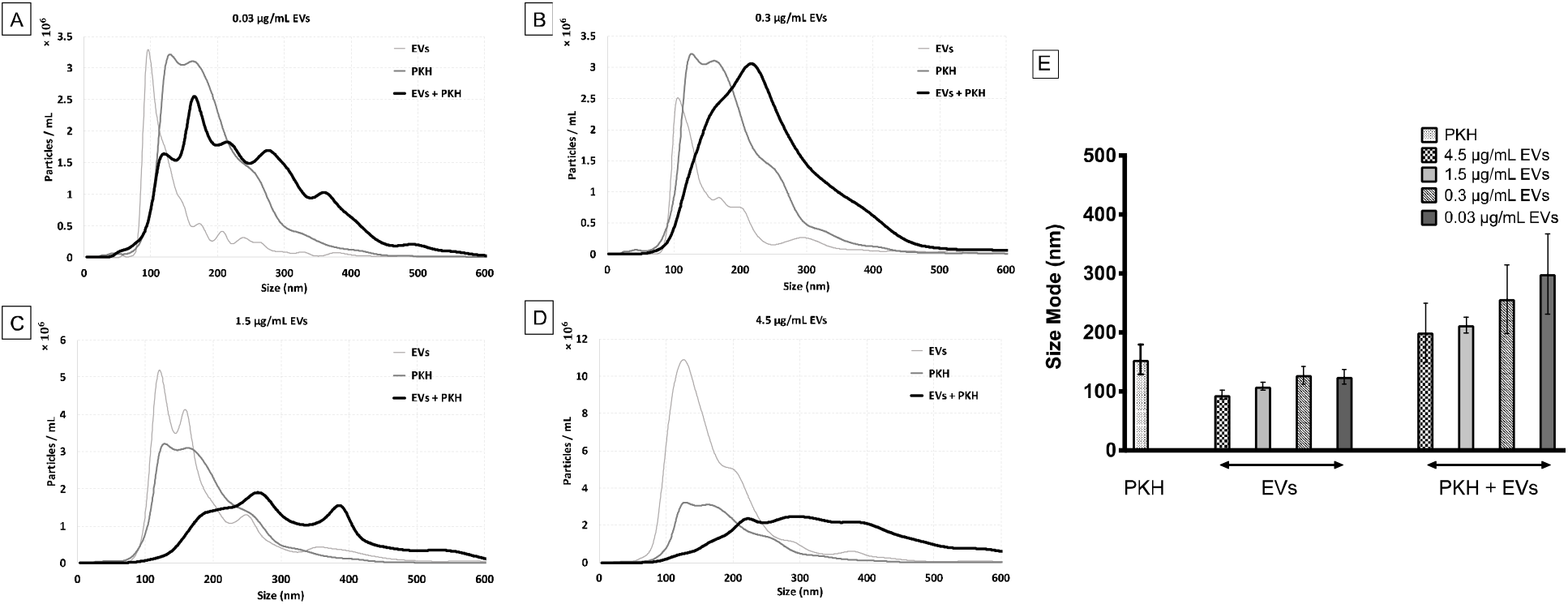
Effect of EV concentration on the size distribution of particles in PKH-labelled EV samples evaluated by Nanoparticles Tracking Analysis (NTA). A-D) Size distribution of PKH-labelled EV samples with different EV concentrations (0.03, 0.3, 1.5, 4.5 μg/mL respectively) with 4 μM of PKH (n=7). E) Size mode of EV only control, PKH only controls, and PKH-labelled EV samples (Error bars represent 95% confidence interval of the mean).

Labeling EV with PKH caused a shift in the size of EVs from ~100 nm for unlabeled EVs to ~200 nm for PKH labelled EVs. Studies have found that using inorganic particles, a similar size shift from 100 nm to 200 nm decreases the cellular uptake efficiency and kinetics. Different types of nanoparticles and cells have been investigated such as mesoporous silica nanoparticles on Hela cells by Lu et al. [35], fluorescent latex beads on B16 cells by Hökstra et al. [42], polystyrene nanoparticles on Caco-2 and MDCK cells by Kulkarni et al. [30]. In addition to cellular uptake efficiency, Hökstra et al. [42] have shown that nanoparticles smaller than 200 nm can be taken up by clathrin-coated pits, while larger particles tend to be internalized by caveolae-mediated processes. This suggests the size shift due to the PKH labelling may change both the cellular uptake level and mechanism of endocytic internalization of PKH labelled EVs.

Furthermore, Kulkarni et al. [30] studied the biodistribution of intravenously injected polystyrene nanoparticles in different rat’s organs. They found that 200 nm nanoparticles showed higher accumulation in both liver and spleen compared to 100 nm nanoparticles. Again, this raises the possibility that increasing the size of EVs by labelling with PKH may affect the EV’s biodistribution. Gangadaran et al. [35], examined the biodistribution of extracellular vesicles (EVs) including EVs using DiR (a similar lipophilic dye as PKH) and found that lipophilic labelling of EVs increased the localization of EVs in liver and spleen. This change in the biodistribution may be due to the increasing in size of EVs after labelling by lipophilic dyes. In summary, the size shift towards larger particles caused by PKH labelling of EVs is likely to change the cellular uptake level and internalization mechanism as well as the biodistribution of EVs, reducing its validity as an EV tracer.

As opposed to lipophilic dyes which may generate larger species, potentially through PKH nanoparticles fusion/aggregation or PKH dyes intercalation with EVs, it is anticipated that direct protein labelling of EVs with fluorescent compounds will not generate these larger EV species. In order to study this hypothesis, the protein binding dye 5-(and-6)-Carboxyfluorescein Diacetate Succinimidyl Ester (CFSE) was used to label EVs. CFSE dyes are membrane permeable chemical compounds, which covalently bind to proteins and fluoresce after ester hydrolysis of the dye in the lumen of the EV [43–46]. Consistent with this hypothesis, Morales-Kastresana et al. [23] and Pospichalova et al. [28] found protein binding fluorescent compounds did not form nanoparticles and did not increase EV size after labelling. It was further shown that unbound CFSE dyes did not interfere with the measurements [28].

EVs were labelled with CFSE dyes using a previously established protocol [23] and the size of particles and fluorescent intensity was explored. Compared to the CFSE and EVs only controls, several brighter features were seen which are likely CFSE-labelled EVs (Figure 5B). Additionally, the line scan of CFSE-labelled EV samples showed spikes in the CFSE-labelled EV sample indicating the presence of CFSE-labelled EVs (Figure 5B). Size distribution and quantitative determination of the size mode showed no significant change in CFSE-labelled EVs compared to unlabelled EVs which was in agreement with findings of Morales-Kastresana et al. [23] and Pospichalova et al. [28]. As opposed to PKH labelling of EV which increases the size, this result suggests that CFSE dye labelling maintains the normal size of EVs which precludes any size related cellular uptake and biodistribution aberrancies. It is important to note that CFSE is not the only alternative to PKH dye. A new class of lipophilic dye (MemBright) also do not form aggregates during labelling of EVs which is essential for tracking EVs [25,26].

**Fig. 5.**
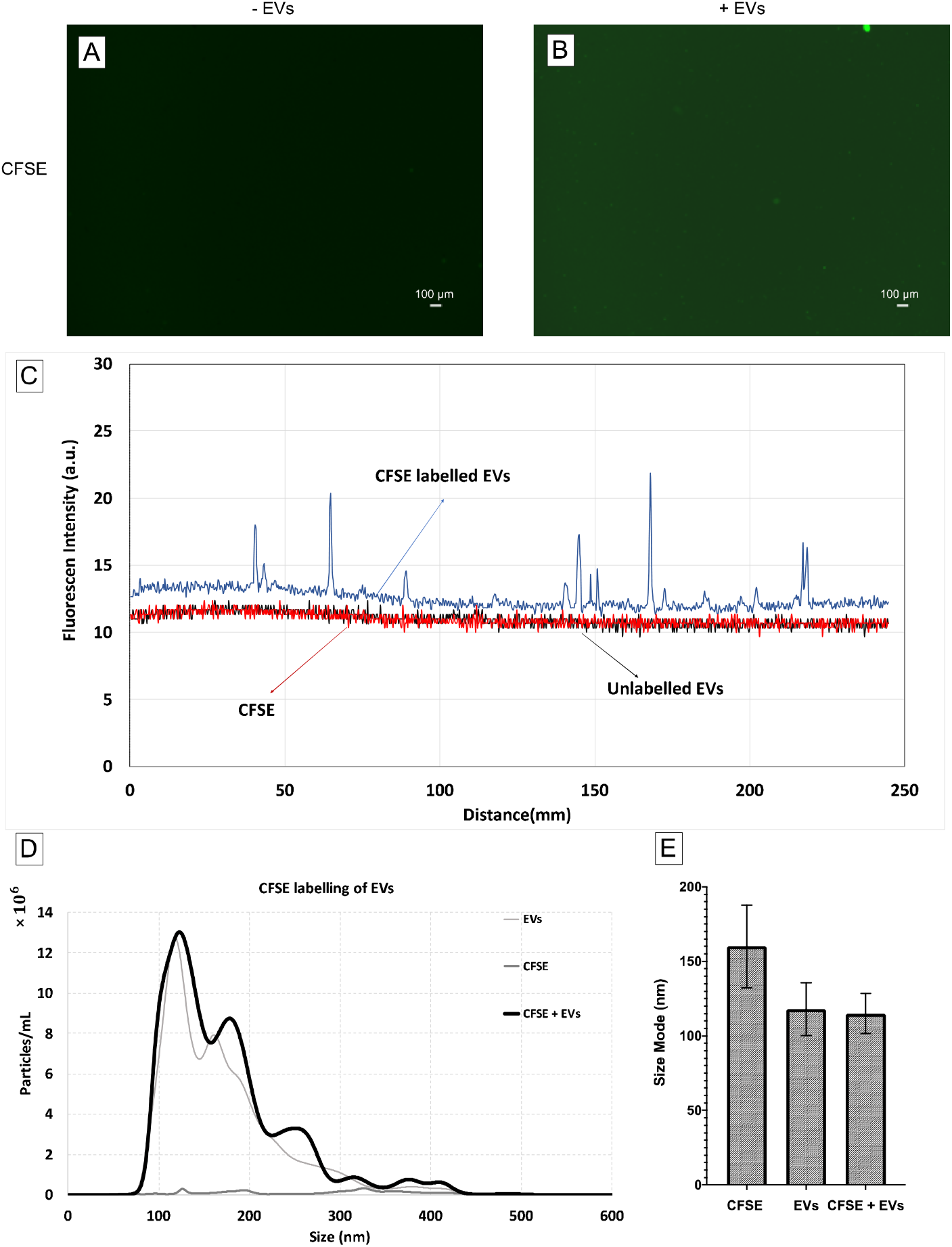
Size characterization of CFSE EV labelling by Nanoparticles Tracking Analysis (NTA). A) Size distribution of EV only control, CFSE only control, and CFSE-labelled EV sample (n=7). B) Size mode of EV only control, CFSE only control, and CFSE-labelled EV sample (Error bars represent 95% confidence interval of the mean).

## Conclusions

Lipophilic labelling of EVs has been shown to have multiple drawbacks. First, non-specific labelling of non-EV biofluid components such as microvesicles, lipoprotein particles, and proteins [22]. Second, the *in vivo* biodistribution of EVs has been shown to be altered by lipophilic labelling [30]. Third, the formation of PKH nanoparticles was observed which can potentially cause false positive results in EV cell uptake and biodistribution studies. Fourth, a size shift in the size of EVs was shown after labelling, most likely due to the PKH nanoparticles fusion/aggregation and PKH dyes intercalation with EVs. These larger species formed after PKH labelling of EVs may cause aberrancies in cellular uptake, biodistribution and half-life circulation.

Here, the relative ratio of PKH to EVs was systematically studied in order to minimize the EV size shift towards larger particles by PKH labelling while maintaining the fluorescent detection of labelled EVs. In all conditions tested, formation of larger species after labelling was detected, even in the conditions where the PKH level is below the fluorescent detection level. In contrast to lipophilic dyes such as PKH, protein binding dyes like CFSE did not cause a size shift in labelled EVs, suggesting that CFSE may be a better labelling option for EVs.

## Acknowledgements

Research reported in this publication was supported by NIGMS of the National Institutes of Health under award number R35GM119623 and NSF award number IIP 1660177 to TRG.

